# SARS2Mutant: SARS-CoV-2 Amino-Acid Mutation Atlas Database

**DOI:** 10.1101/2022.06.12.495856

**Authors:** Karim Rahimian, Mohammadamin Mahmanzar, Bahar Mahdavi, Ehsan Arefian, Donna Lee Kuehu, Youping Deng

## Abstract

The coronavirus disease 19 (COVID-19) is a highly pathogenic viral infection of the novel severe acute respiratory syndrome coronavirus 2 (SARS-CoV-2), resulting in the global pandemic of 2020.A lack of therapeutic and preventive approaches including drugs and vaccines, has quickly posed significant threats to world health. A comprehensive understanding of the evolution and natural selection of SARS-CoV-2 against the host interaction and symptoms at the phenotype level could impact the candidate’s strategies for the fight against this virus.

<u>SARS</u>-CoV-<u>2 Mutation</u> (SARS2Mutant, http://sars2mutant.com/) is a database thatprovides comprehensive analysis results based on tens of thousands of high-coverage and high-quality SARS-CoV-2 complete protein sequences. The structure of this database is designed to allow the users to search for the three different strategies among amino acid substitution mutations based on gene name, geographical zone or comparative analysis. Based on each strategy, five data types are available to the user: mutated sample frequencies, heat map of the mutated amino acid positions, timeline trend for mutation survivals and natural selections, and charts of changed amino acids and their frequencies. Due to the increase of virus protein sequence samples published daily showing the latest trends of current results, all sequences in the database are reanalyzed and updated monthly. The SARS-2Mutant database providescurrent analysis and updated data of mutation patterns and conserved regions, helpful in developing and designing targeted vaccines, primers and drug discoveries.

## INTRODUCTION

The new subfamily member of *Coronavirinae,* subsequently named severe acute respiratory syndrome coronavirus 2 (SARS-CoV-2) caused coronavirus disease 2019 (COVID-19), which appeared for the first time in the Wuhan State of Hubei Province in China, in early December 2019 [1, 2]. With the worldwide spread of SARS-CoV-2, large populations have been infected, which already accounts for more than6.1 million deaths and about 493 million cumulative cases globally, as of 8 April 2022 (WHO, Coronavirus (COVID-19) Dashboard, covid19.who.int). In addition, studies indicate that the numbers of indirect covid-19 deaths, such as heart disease and stroke, increased rapidly on a daily basis [3, 4], prompting attention to this disease which has become one of the major treatment priorities of all countries and the World Health Organization (WHO) [2]. SARS-CoV-2 is the seventh coronavirus known to infect humans [5] and is classified as a *Sarbecovirus* subgenus, *Betacoronavirus* genus, and *Orthocoronavirus* subfamily member belonging to the *Coronaviridae* family [6]. High throughput data techniques such as Next Generation Sequencing (NGS) revealed that this virus derives about 80% of its genome sequence identity from the severe acute respiratory syndrome coronavirus (SARS-CoV), which emerged in 2002-2003 [7]. As of April 2022, almost 10 million full genomes are available via the Global Initiative on Sharing All Influenza Data (GISAID), which is one of the main pandemic genome databases.

The SARS-CoV-2 is 50-200 nm in diameter, has a lipid-enveloped, a positive sense, and is a single-stranded RNA virus. The full-length RNA genome is comprised of 29,903 nucleotides (nt) consisting of the open reading frames 1a and 1b (ORF1ab). ORF1ab encodes ORF 1 polyproteins that are proteolytically processed into 16 mature non-structural proteins (NSPs) that play critical roles as regulatory proteins in viral RNA replication and transcription. Moreover, SARS-CoV-2 contains genes that encode four major structural proteins, including spike surface glycoprotein (S), an envelope protein (E), membrane glycoprotein (M), and nucleocapsid phosphoprotein (N), all of which are responsible for the infectious virion assembly. The N protein packages the RNA genome into a helical ribonucleocapsid. The S, E, and M proteins generate the viral envelope (Figure. 1). The S protein is comprised of two functional subunits which are involved in viral interactions with a host cell receptor angiotensin-converting enzyme 2 (ACE2) (S1 subunit), as well as mediating the fusion of the host and the viral membrane (S2 subunit). Hence, it is considered as a potential therapeutic target for antiviral drug development. Interspersed between structural genes are several other genes which are translated to proteins called accessory factors. Although the biological functions for each of these genes has been determined, a number remains unclear [7–9].

**Figure 1.**
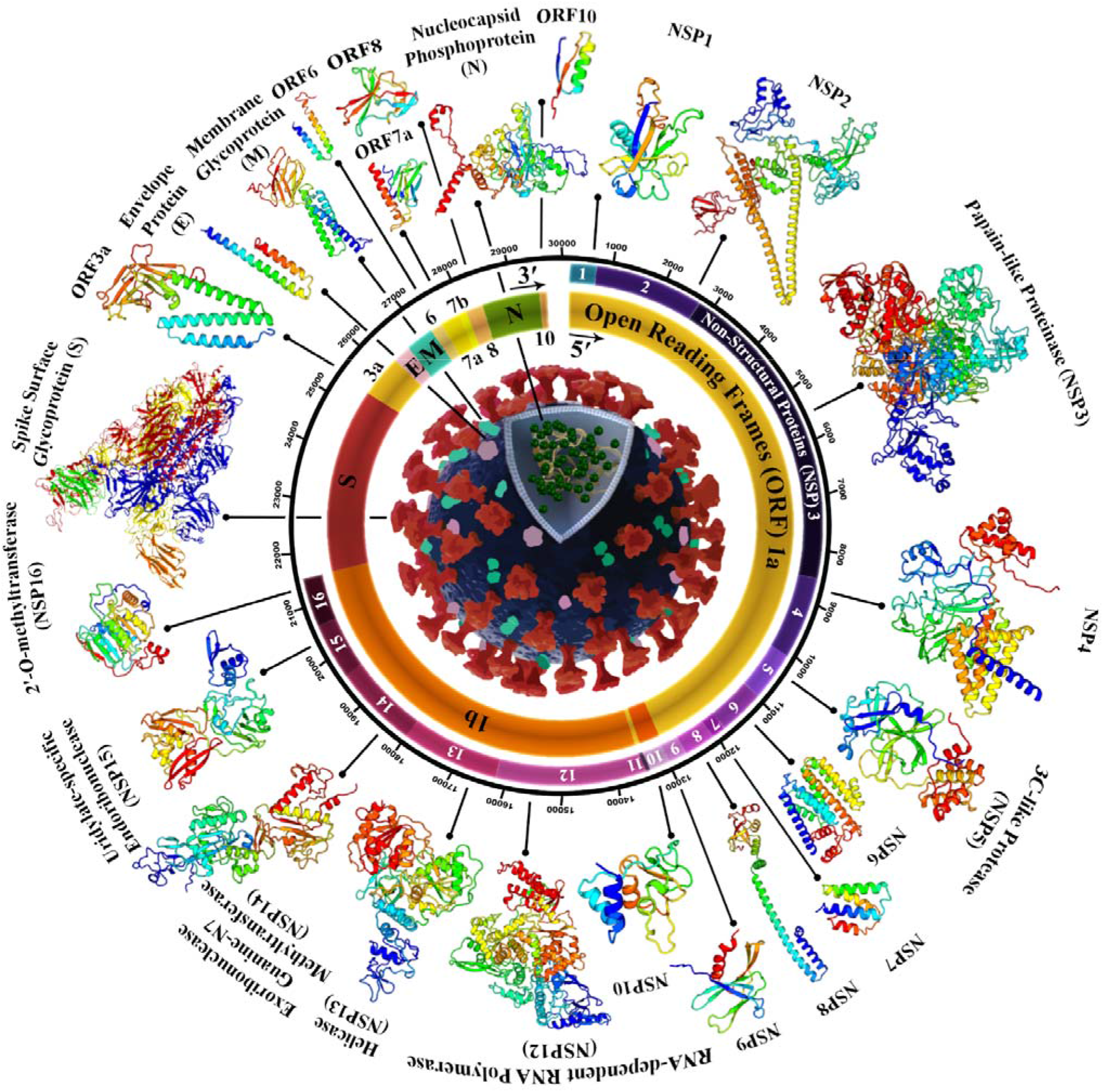
Schematic view of the SARS-CoV-2 particles, genome arrangement, and proteome organization. SARS-CoV-2 is an enveloped positive-sense single-stranded RNA betacoronavirus with a ~30kb polycistronic genome that encodes non-structural proteins (ORF1a and ORF1b, that are processed into Nsp1-16) at the 5’-end, and structural proteins (S, E, M, and N), and several other accessory factors (ORF3a, 6, 7a, 7b, 8, and 10) at the 3’-end. 3D structural models of protein are obtained from I-TASSER [18].

During the replication process, all viruses, including SARS-CoV-2, mutate. This phenomenon includes the occurrence of occasional mistakes during replication inside host cells. Most of the time, mutations make no alterations to the function of the virus, and it may even result in weakening it. However, sometimes they accumulate advantages such as boosting infection ability, evading the immune system, and expanding within the population, resulting in a variant designation[10–12].

The spread of different variants of coronavirus is of great importance. For instance, genetic variants may interfere with diagnostic tests and cause false-negative molecular detection, elevating SARS-CoV-2 spreading potential in the presence of antibodies[11]. Additionally, they can also diminish the efficiency of therapeutical approaches or even increase the severity of COVID-19. To date, the most effective strategy against the coronavirus epidemic is universal vaccination. One of the notable reasons concerning the emergence of new variants is the probability of reinfecting individuals or those fully vaccinated (vaccine breakthrough) [12].

Due to the importance of SARS-CoV-2 amino acid mutations and their correlations to the available therapeutic methods, public access for governments and health institutions monitoring mutations are essential during the evolution of the virus.. On the other hand, the launch and availability of the desired SARS-CoV-2 single amino acid variations (SAVs) will also greatly help healthcare professionals to prescribe the proper evidence-based medication [13, 14]. In addition, access to these databases is of great help to researchers and scientists working to develop new molecular diagnostic techniques, vaccines, and therapies for COVID-19.

So far, a limited number of databases have been established for the SARS-CoV-2, but mainly to classify viruses at the genome mutations and evolutionary level[15–17]. However, due to the importance of mutations at the amino acids level and the mutation effects on protein function, a database that classify genes and reported mutations at the protein level based on continent, country, and timeline has not yet been available until now. The Sars2Mutant database analyzes and identifies mutations at the protein level and positions, reports the exact loci of the mutations, classifies the modifications based on the frequency in each gene, and identifies geographical hotspot regions and those highly conserved regions aligned to the reference Sequence., Other important features of the Sars2Mutant database are ease of use, transparency of data presentation, and user friendly experience. Researchers can use the database quick access module to study SARS-CoV-2 mutations by genes and geographical zone or compare mutation frequency status across continents and countries.

## MATERIALS AND METHODS

### Data collection

The first version of the SARS2Mutant database contains 4 million high-quality and high-coverage SARS-CoV-2/hCoV-19 protein sequences downloaded from Global Initiative on Sharing Avian Influenza Data (GISAID, https://www.gisaid.org/) [19–21] from November 2019 until June 2021.

### Pre-processing and quality control

Non-human samples (such as bat and pangolin), those with less or more than reference length amino-acid sequences (AAs), samples containing non-specified AAs (reported as X), and no reported geographical location were omitted. Ultimately, on 28 April 2022, more than 10.5 million samples were included in this study. The whole process was performed by applying python libraries such as ‘Numpy’ and ‘Pandas.’. This included samples summarized in Table1 which followed a step by step filtering process.

**Table 1.** The number of included samples which sorted step by step.

| Gene name | Total | Criteria trimming |  |  | Remained |
| --- | --- | --- | --- | --- | --- |
|  |  | Not match length | Sequence contain X | Non human |  |
| NSP1 | 10,261,296 | 586,827 | 433,559 | 7,298 | 9,233,612 |
| NSP2 | 10,280,285 | 23,067 | 940,384 | 5,712 | 9,311,122 |
| NSP3 | 10,285,623 | 2,240,262 | 1,971,864 | 3,322 | 6,070,,175 |
| NSP4 | 10,283,271 | 18,737 | 1,357,844 | 7,135 | 8,899,555 |
| NSP5 | 10,283,326 | 13,584 | 650,306 | 7,390 | 9,612,046 |
| NSP6 | 10,283,424 | 4,573,308 | 556,800 | 4,278 | 5,149,038 |
| NSP7 | 10,283,211 | 1,831 | 113,086 | 7,955 | 10,160,339 |
| NSP8 | 10,283,099 | 2,993 | 321,269 | 7,907 | 9,950,930 |
| NSP9 | 10,283,166 | 2,303 | 171,536 | 7,942 | 10,101,385 |
| NSP10 | 10,283,220 | 5,178 | 386,807 | 7,783 | 9,883,452 |
| NSP11 | 10,282,621 | 406 | 248,550 | 8,011 | 10,025,654 |
| NSP12 | 10,283,038 | 15,751 | 1,417,103 | 6,870 | 8,843,314 |
| NSP13 | 10,283,161 | 8,417 | 947,020 | 7,320 | 9,320,404 |
| NSP14 | 10,283,,125 | 16,466 | 1,963,818 | 6,608 | 8,296,233 |
| NSP15 | 10,283,011 | 11,275 | 1,082,988 | 5,821 | 9,182,927 |
| NSP16 | 10,284,335 | 14,175 | 987,187 | 6,527 | 9,276,446 |
| ORF3a | 10,285,527 | 218,561 | 582,697 | 7,266 | 9,477,003 |
| ORF6 | 10,282,433 | 142,987 | 132,233 | 7,818 | 9,999,395 |
| ORF7a | 10,282,309 | 732,602 | 496,921 | 7,595 | 9,045,191 |
| ORF7b | 10,269,432 | 477,420 | 491,389 | 7,753 | 9,292,870 |
| ORF8 | 10,278,801 | 5,582,221 | 495,547 | 4,480 | 4,196,553 |
| ORF9b | 10,282,579 | 3,215,630 | 227,157 | 5,499 | 6,834,293 |
| ORF9c | 10,280,158 | 485,696 | 156,699 | 7,228 | 9,630,535 |
| S | 10,391,577 | 8,545,727 | 894,375 | 1,015 | 9,504,60 |
| E | 10,284,323 | 1,97,859 | 163,922 | 8,012 | 9,914,530 |
| M | 10,284,401 | 94,574 | 1,322,197 | 7,166 | 8,860,464 |
| N | 10,283,011 | 3,265,363 | 647,251 | 4,939 | 6,365,458 |

### Variants calling and functional annotation

Sequence alignments were made by detecting total single amino-acid (AA) variations against the reference genome, Wuhan-WIV04 (EPI_ISL_402124). Wuhan-WIV04 genome is the fulllength protein sequence reference of the SARS-CoV-2 identified from China in December 2019, known as the reference sequence in GISAID that determined each gene location precisely. Another reference sequence reported in NCBI is Wuhan-WIV04 (NCBI: NC 045512.2), which is one of these two sequences, Wuhan-WIV04 and Wuhan-Hu1, and are the same in protein levels. At the genome levels, Wuhan-Hu-1 has 12 more polyAs at the end of the RNA genome, but protein levels are not affected. All SARS-CoV-2 sequence mutations is scheduled to be updated monthly, and powered by the GISAID database.

The complete reference sequences of SARS-CoV-2 were captured from the GISAID database. Access to this database is by permission of John A. Burns School of Medicine Department of QuantitativeHealth Sciences, and data preprocessed with Python libraries. After filtering low-quality samples and removing white spaces within the series, we designed a unique library based on an exact match algorithm pairwise to align SARS-CoV-2 sequences with the reference genome Wuhan-WIV04. This library is available on https://github.com/sars2mutant/covid_db, which can handle the big data sequence pairwise aligner that aligns long protein sequences in the FASTA format based on our strategy for analyzing whole SARS-CoV-2 sequences. The pairwise alignment results are not affected by other sequences. Each sequence was aligned separately with the reference genome, and the variations were reported. The worldwide mutation rate for each gene was obtained by dividing the number of identified mutations by the number of mentioned gene samples.

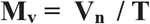

Where the M_v_ is the mutant variations, V_n_ is the mutated amino acid number, and T is the total number of included samples. To report the rate of mutations for each continent, the same method was used, dividing the total mutation number of the specific gene by the number of the mentioned gene,

The current database is specialized for introducing SARS-CoV-2 protein mutations, the details of each mutation including the exact location, its geographical incidence, and the replaced AA. Therefore, each sequence is labeled and an information profile for classification data is created. The data structure can search and categorize data based on the mutant ratio in each part of the protein sequence (nsp1…nsp16, S, E, M, N, ORFs), the concurrence, and mutation frequency by geographical zone.

### Platform architectural design and structure

The SARS2Mutant web platform and relational database connection were implemented using Django package in Python 3.9.7 programming environment for the backend. HTML, CLS, and Javascript were used to design the frontend. An object-oriented architecture was designed and implemented in a relational database (MySQL) to store the annotated variants instead of the conventional spreadsheet file (CSV/VCF) to allow further flexibility when formulating search queries and alleviate database load by reducing data duplication that causes reduced data load time. The database architecture and relationships between tables are shown in Figure 2.

**Figure 2.**
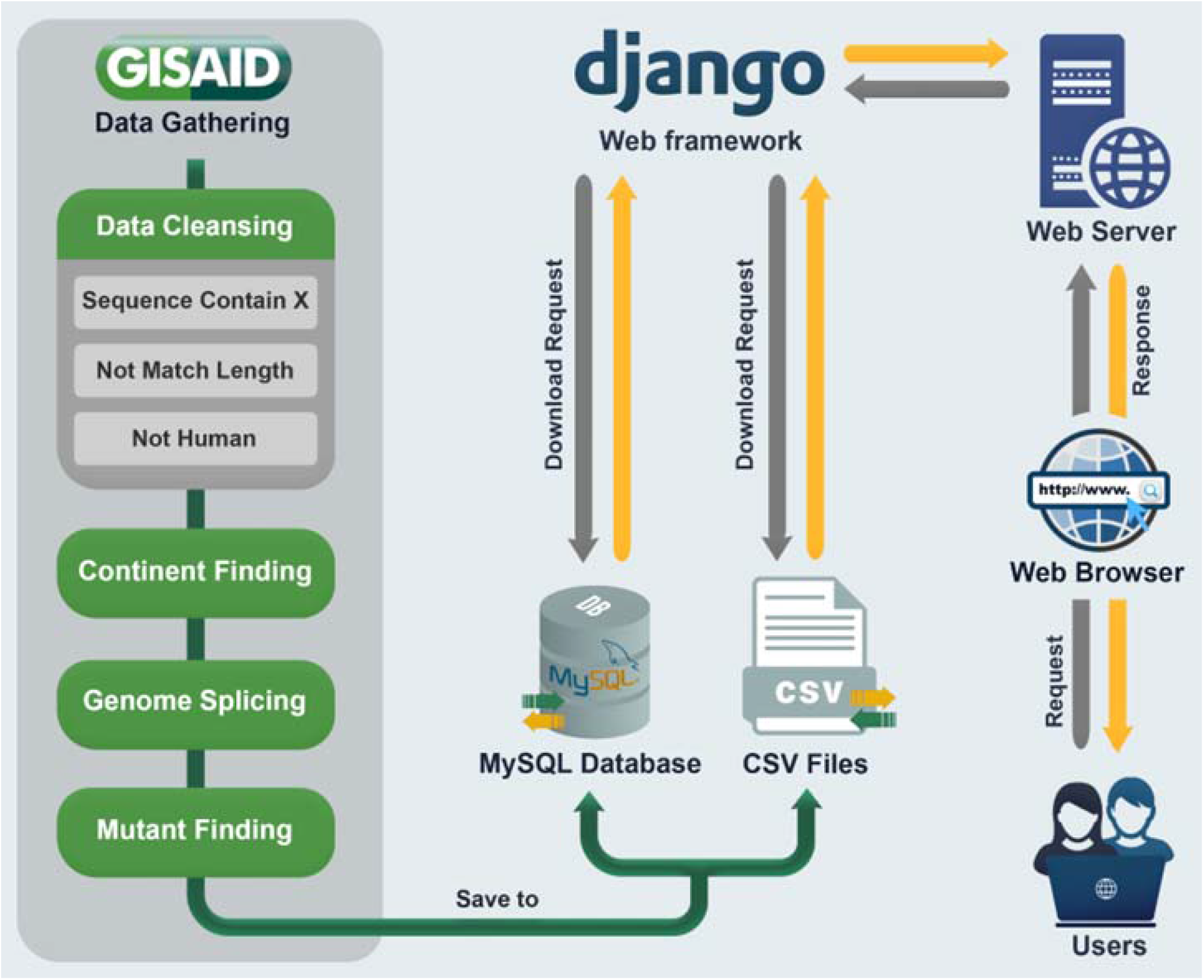
Database architecture and ork flow. Request mechanisms from user to database.

## RESULTS

The home page of the SARS2Mutant database provides a summary of our values and visions regarding site design (Figure 3A). To assist the researcher in formulating a research idea, a table of the SARS-CoV-2 genes have been designed and their function summarized clearly to facilitate with the interpretation of the results.

**Figure 3.**
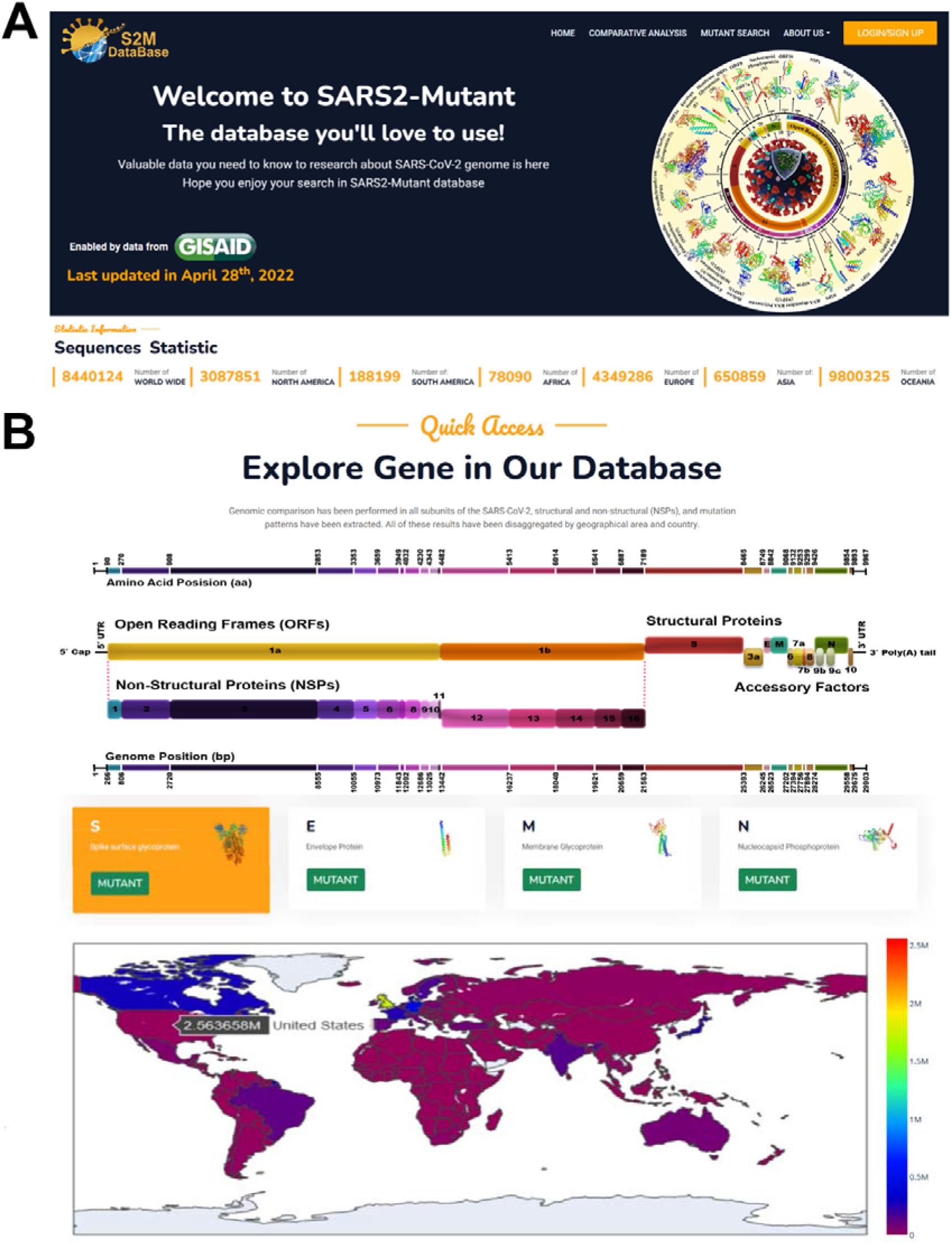
Database structure. (A) showed the home page, services and facilities. (B) represent quick access part of database that helps the user find the results quickly and clearly.

Sars2Mutant taskbar includes a set of ‘Home,’ ‘Comparative Analysis’, ‘Mutant Search,’ ‘About Team’, and ‘Login / Sign Up’. Tabs 1, 2 and 3 represent_three main search strategies: ‘ Quick access’, ‘ Comparative analysis’ and ‘ Mutant Search’ provided in this database.

At the top of the site, a section called “Quick Access” includes 28 cards which provides quick access links. By placing the mouse on each card, the analysis selected by the user transfers into the results page providing two categories for review:

1. The top 100 mutation reports includes mutation occurence distributions, AAs mutation position, substitution AA, hotspot map, and top 10 sustainable monthly tracking timeline). To access total data, the user must make register for a free account which allows a download of CSV files from each analysis.
2. Top mutation selections are based on the occurrence frequency from different geographical locations mapped on a worldwide graph.

The “quick access” section allows users to access the results faster and easier (Figure. 3B).

At the bottom of the home pageis an interactive ‘Worldwide Mutation Distribution’. This section allows users to find more categorized information about mutation frequency in the countries/areas by moving the mouse over the maps to any part of their area of interest. A table within the ‘Worldwide Mutation Distribution’ tab can be downloaded showing the counts of SAVs in each country/area.

The details of each search strategy and outputs provided by the current version of SARS2Mutantare provided.

### Protein name search, “Quick Access”

The ‘*Protein Name search*’ allows users to explore SAVs in a particular region of the viral-specific protein (‘*Protein-based search*’). SAVs within the selected protein is presented in the pie chart, Heatmap, Stacked plot, Timeline chart and worldwide map, which shows mutation frequency among samples, the comparative analysis between hotspot versus conserved region in the referenced gene, substitution AAs name and frequency, monthly mutation survival trend and geographical mutations distribution.

For example, to find the spike protein (S) mutations, in part A, there is a description of protein activity and its official sequence (Figure 4A). The part B pie chart shows that 4.82%, 26.31%, 25.31%, 13.75%, and 29.79% of samples are conserved, have one mutation, have two mutations, and have three mutations and more than four mutations, respectively, worldwide (Figure 4B). In the heatmap, the x-axis represents the hotspot positions, and the total length of each gene is divided into 10 sections. In the y-axis, it expresses the mutation frequency of each position. In the spike example, 508 – 635 AA positions are the areas with the most frequent mutations in the gene (Figure 4C). To track the type of AA changes and their frequency in each point position, the y-axis of the stacked plot shows the Log frequency for comparison. On the x-axis is the name of wild-type AAs and their position. For example, in the case of spike protein, D614 was detected as high point AAs that substitute Glycine (D614G) with high frequency, and the second highest substitution is Asparagine (D614N) (Figure 4E). Each mutation based on evolutionary parameters during the pandemic showed different survival patterns. The timeline chart shows the sustainability of each mutation during the month. The y-axis showed the mutation rate, and the x-axis represents each month.

**Figure 4.**
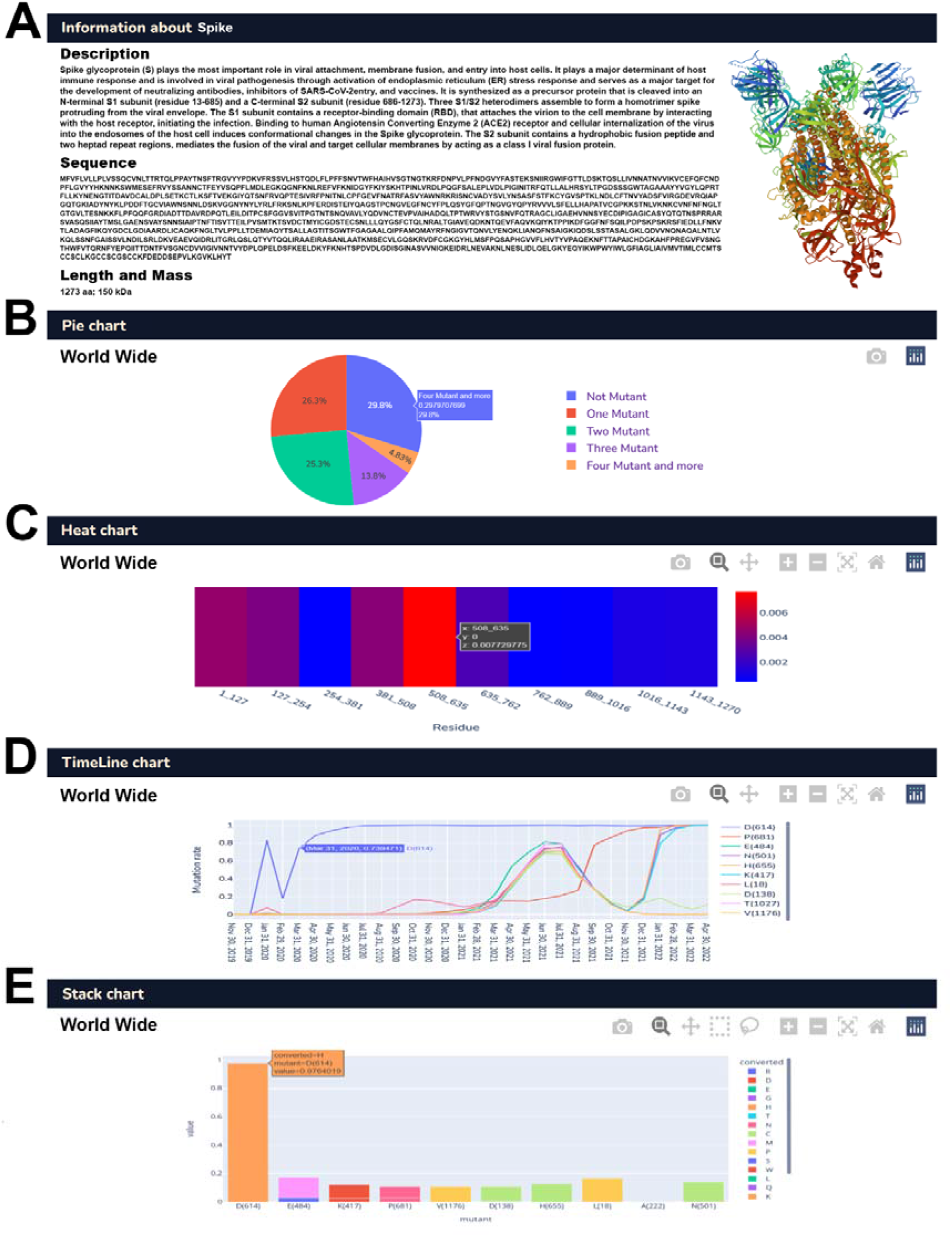
Database data representation structure. (A) Protein description, functions and official sequence. (B) mutation frequency among all analyzed samples. (C) Hotspot genome positions versus conserved. (D) Mutation detection trend during the month. € Mutation frequencies and Substitution amino acids.

The color of the line represents the mutation point position of mutations. It could indicate which positions could play an essential role in that specific gene function, which could be more adaptable to different situations. Based on our outcomes, the chart shows the D614 position mutation was observed for the first time in Dec 2019, and after that in Feb 2020, where the increase in mutation numbers began. The second top mutation frequency, E484 position, shows an uptrend from Nov 2021 to the present (Figure 4D). In the last part, the worldwide map was designed to represent the distribution of the specific mutations globally.

It’s notable that in all the mentioned graphs and maps, users are allowed to find more categorized information about mutation frequency in the countries/areas by moving the mouse over the maps to any part of their specific interest or by free registration for database access to the total data, representing more than 100mutations.

### SAVs birth query, “Mutant Search”

Similar to other viruses, SARS-CoV-2 has created genetic diversity via temporally accumulated mutations. The ‘SAV birth query’ is set up for users to overview the geographical zones where SAVs are discovered, and for the exact AA position and substitution AA. This means that the area here the mutation were observed is shown by selecting particular mutations. (Figure 5A).

**Figure 5.**
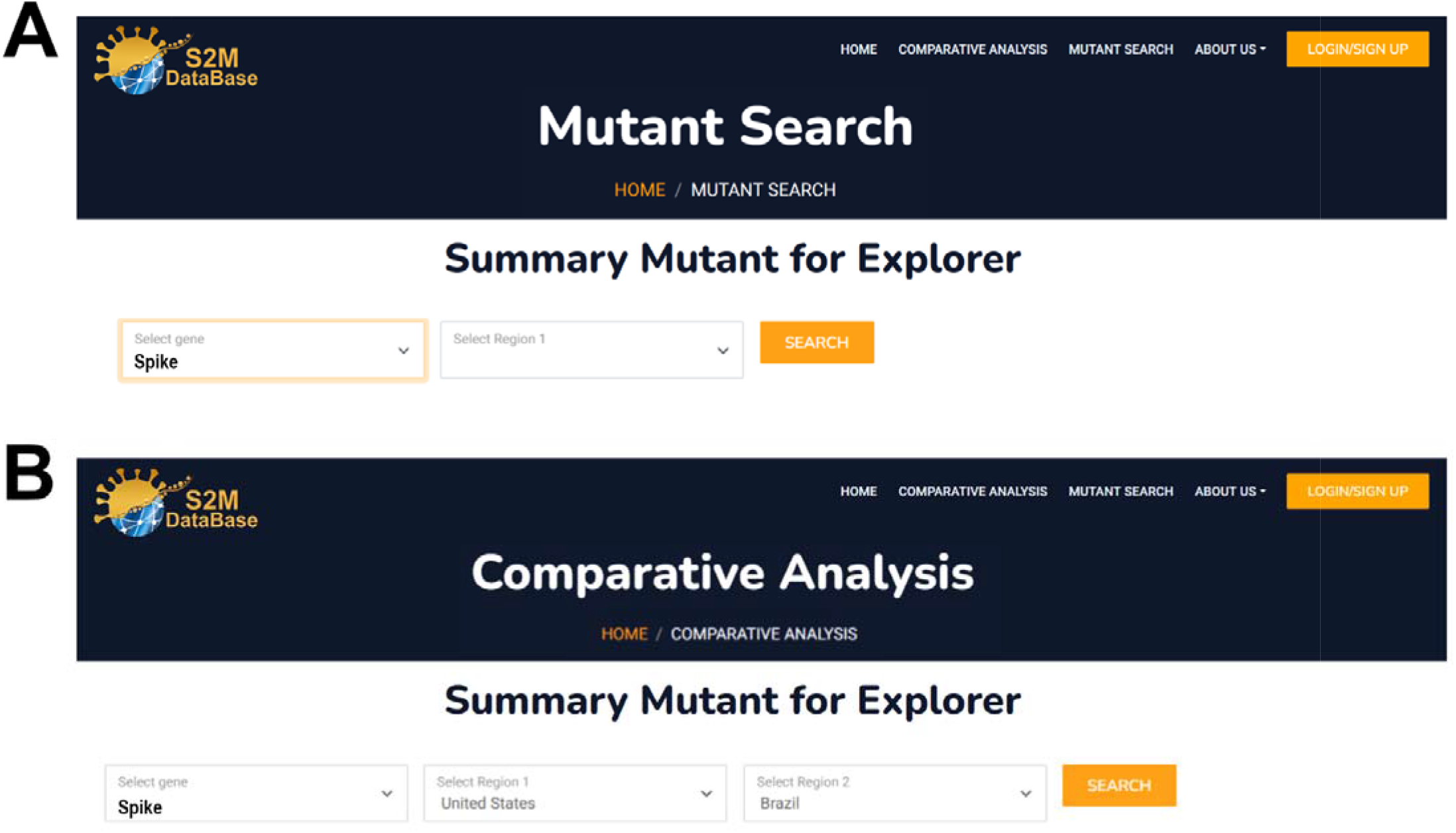
Database search strategy. (A) allow the user to search based on the candidate gene. (B) Users can search and compare the candidate gene between two different areas.

### Search mutations based on Geographical Zone, “Comparative analysis”

The ‘Continent/ Country search’ helps users focus on SAVs identified in a specific continent or Country (‘Region-based search’). The summary of SAVs in the area includes names and positions of SAVs, numbers, and percentages of viral genomes carrying the SAV and replaced AA reported in the selected area. Moreover, users can also filter or do a quick search on SAVs given attractive attributes by adding the keyword in the search box. Users can also select the particular gene and the target country/continent to compare the results between the two regions. All the results presentation structures are the same as the “Quick Search” section and SAVs within selected areas are presented in the same graphs. All data is available to download through CSV file format (Figure.5B).

## DISCUSSION

Numerous online databases of SARS-CoV-2 mutations have been developed over the last two years [24–29]. In comparison, SARS2Mutant database provides a user-friendly environment for easy operations to obtain a holistic overview of SARS-CoV-2 SAVs. The remarkable difference between the SARS2Mutant database and similar available databases such as GESS [29], Covariants [17], COVMT[15], VirusViz[16] and IDBSV [30] is simply that data visualization over thousands of annotated records cover complete SARS-CoV-2 genome and other numerous parameters summarized and compared in Table 2.

**Table 2.**
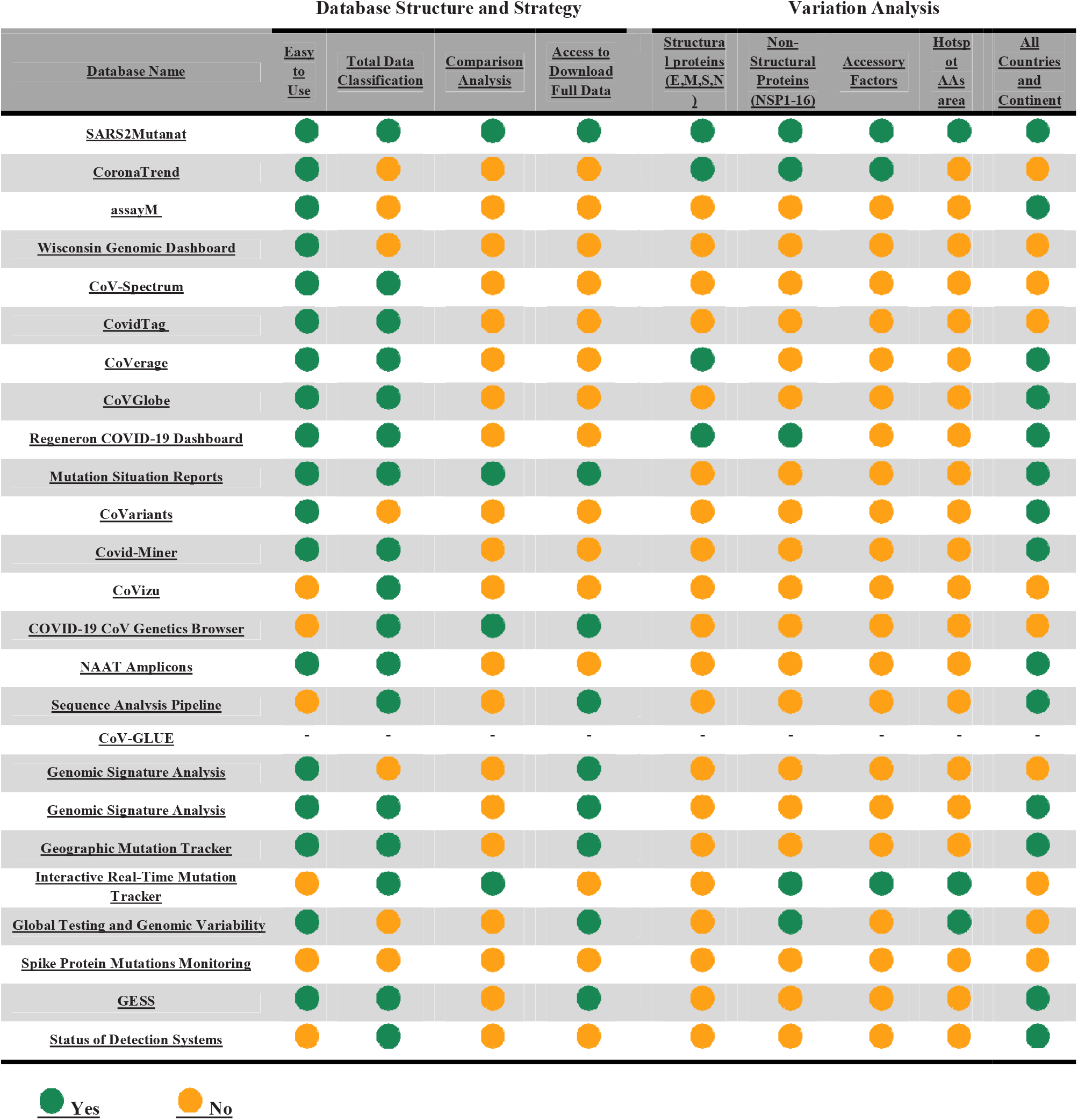
Comparison checklist between same concept database with SARS2mutant.

Each protein has a specific profile and includes classified data that can be compared with other proteins. Additionally, according to a study on the protein level, only important mutations which appear at the phenotype level are reported, and silent mutations which do not result in a change in the AAS and subsequently have no effect on protein folding and function are not considered in the reports. In addition, according to the SARS-CoV-2 gene structure, the exact location of mutations in each gene was reported separately by the SARS2Mutant database, which will be very effective in investigating for drug and vaccine effects. Researchers could search for specific mutations based on proteins, AA location and selective substitution AA.

Since proteins play a role in the presence of phenotypes, the accumulation of various mutations at the level of AAs can change the structure of the proteins and ultimately produce a new phenotype characterization that can give a new feature to the mutant virus or alter the impact on its pathogenesis and virulence [31, 32].

The evolution of viruses depends on the co-occurrence of various mutations in multiple genes or within a single gene. The accumulation of mutations in viruses has resulted in drugresistant or vaccine escape mutants, leading to a continuous need to design new drugs or vaccines [33]. Ongoing screening of functional mutations might help provide insight into the evolution and genetic diversity of SARS-CoV-2, which is also critical for developing efficient antiviral drugs or vaccines against this virus [34]. Vaccination, antiCoV therapeutic strategies, and diagnostic products are the main substantial tools for inhibiting global disease. Extensive global research and effort have been dedicated to developing an efficient vaccine to elicit a rapid and robust immune-protective response and provide a strong and long-lasting immunity against different strains of the SARS-CoV-2 [35],

On the one hand, the ideal vaccine must have strong immunogenicity. It should be able to elicit targeted humoral and cellular immune responses via specific epitopes. On the other hand, it should also be less immunotoxic and non-allergenic [35–37]. Moreover, to provide a universal antibody to neutralize different viral strains, the conservation of targeted sequences such as the T and B cell epitopes that can elicit cellular and humoral immune responses must also be considered. The SARS2Mutant database can be of great help to calibrate mutation rates for specially designed regions.

There exists other functional mutations which can give rise to drug resistance. Therefore, screening the mutations to understand and manage the mechanisms of SARS-CoV-2 drug resistance is imperative to find viable drug target candidates for designing effective antiviral drugs [38–40]. Also, functional mutations can impact the results of diagnostic tests and lead to false-negative using rapid antigen tests. So it is essential to monitor the SARS-CoV-2 mutations to develop stable and reliable diagnostic tests [41–43]. Accordingly, mutation screening investigations in a fast, reliable, and cost-effective way with the help of databases will help develop effective coping strategies during the COVID-19 pandemic.

According to GISAID powered database that used the same data representing different aspects of data, we summarized the difference between SARS2Mutant database and others in Figure.4. Based on two main parts of the database, technical structures and data representations, are evaluated and compare. Although SARS2Mutant database primary goal is to present molecular details, it also shows the epidemiological status, summarized in Figure.4 as the same idea database.

The home page of SARS2Mutant contains information about the number of samples collected.

Search functions on SARS2Mutant offer users more considerable flexibility to browse and search SAV patterns from different aspects, such as protein locations, sample geographical zone, and mutation frequency rate, while focusing on SAV characteristics. Notably, a critical feature of SARS2Mutant is the usage of correlation function on SAVs, where a parameter, concurrence ratio R, is adopted to identify SAVs that occurred simultaneously. SARS2Mutant database also provides a novel process for SAV birth query to monitor newly occurred SAVs each month. Through visualized distributions and graphical diagrams, it helps users better understand the migration, transmission, spread, and evolution trend of SARS-CoV-2.

The goal of SARS2Mutant is to provide a user-friendly database to explore the associations and interactions between SAVs. In general, by fetching the data of SAVs and using functions embedded in SARS2Mutant to analyze their significant features, users may gain new insights into the molecular drivers of SARS-CoV-2 transmission, migration, and evolution.

In the hope of creating public safety in countries and overcoming this virus, we intend to increase the number of sequences of the virus sequenced for each country, to enclose the distribution of this data to countries. So that appropriate treatment and vaccination patterns can be suggested for each country tailored to the genetics and blueprints of that region.

Up until 28 April 2022, SARS2Mutant hosted over 60 thousand variations extracted from the analysis of over 10.5 million SARS-CoV-2 samples. Notably, the analysis of these mutations revealed consistent results with the findings from specialized studies and literature.

## Data Availability Statement

GISAID data provided on SARS2Mutant database are subject to GISAID’S terms and conditions (https://www.gisaid.org/registration/terms-of-use/).

All used library and alinger codes are availbe on https://github.com/sars2mutant/covid_db

## Supplementary Data

## Acknowledgments

We sincerely thank all worldwide contributors who have sequenced and shared their data about SARS-CoV-2 in the GISAID database. All data authors can be contacted directly *via* www.gisaid.org.

